# A neurocognitive investigation of the impact of socialising with a robot on empathy for pain

**DOI:** 10.1101/470534

**Authors:** Emily S. Cross, Katie A. Riddoch, Jaydan Pratts, Simon Titone, Bishakha Chaudhury, Ruud Hortensius

## Abstract

To what extent can humans form social relationships with robots? In the present study, we combined functional neuroimaging with a robot socialising intervention to probe the flexibility of empathy, a core component of social relationships, toward robots. Twenty-six individuals underwent identical fMRI sessions before and after being issued a social robot to take home and interact with over the course of a week. While undergoing fMRI, participants observed videos of a human actor or a robot experiencing pain or pleasure in response to electrical stimulation. Repetition suppression of activity in the pain network, a collection of brain regions associated with empathy and emotional responding, was measured to test whether socialising with a social robot leads to greater overlap in neural mechanisms when observing human and robotic agents experiencing pain or pleasure. In contrast to our hypothesis, functional region-of-interest analyses revealed no change in neural overlap for agents after the socialising intervention. Similarly, no increase in activation when observing a robot experiencing pain emerged post-socialising. Whole-brain analysis showed that, before the socialising intervention, superior parietal and early visual regions are sensitive to novel agents, while after socialising, medial temporal regions show agent sensitivity. A region of the inferior parietal lobule was sensitive to novel emotions, but only during the pre-socialising scan session. Together, these findings suggest that a short socialisation intervention with a social robot does not lead to discernible differences in empathy toward the robot, as measured by behavioural or brain responses. We discuss the extent to which longer term socialisation with robots might shape social cognitive processes and ultimately our relationships with these machines.

## 1. Introduction

While any robot we encounter in the real world might not yet possess the emotional intelligence of fictional robots such as C3PO or Wall-E, roboticists are making significant strides in developing robots that can form strong social bonds with humans, thus enabling them to provide long-term support and possibly even companionship to people as a result [1,2]. This interest in developing artificial agents to serve as social companions or assistants has inspired a growing number of studies to examine the extent to which, and under which circumstances, we might engage with ‘social’ robots as we would another person or pet [3,4]. One promising avenue for examining social engagement with another agent is to measure if and how we empathise with that agent when we observe it in apparent pain. Empathy, the observer’s perception and reaction (e.g. sympathy, personal distress) to the emotional state (e.g. pain, distress) of another individual, is a crucial building block of social interactions and relationships [5,6]. Empathy fosters prosocial behaviour, and research examining how we empathise with other people in pain suggests that brain regions engaged when an individual experiences nociceptive pain (direct nerve damage) are similar to those engaged during empathic pain (observing another individual in pain) [7,8]. But how flexible are the behavioural and brain mechanisms underpinning empathy? In the present study, we examine the extent to which humans experience empathic pain when watching a robot “in pain”, and if repeated interaction with this robot, during which deeper social bonds might develop, influences empathic responses to seeing the robot in pain.

A major focus of research and development in social robotics currently is the design and implementation of increasingly sophisticated emotion models in robotic systems [9,10], as well as improved means of communicating emotion by the robots themselves [11-13]. However, as several research groups working at the intersection of robotics and psychology point out, how people respond to emotions expressed by robots in general, and the impact of these emotions on peoples’ emotions in particular, has received considerably less research attention [3,14]. In an attempt to advance understanding of how people might emotionally respond to robots, Rosenthal-von der Pütten and colleagues [15,16] conducted two studies wherein they measured participants’ emotional responses when viewing videos depicting physical violence directed toward a person or a robot (in this case, Ugobe’s Pleo robot, an entertainment robot in the shape of a baby dinosaur that is roughly the size of a small dog). Self-reported data indicated that participants felt an emotional response and empathy when observing a robot in pain. These findings were supported by measures of physiological arousal (skin conductance; [15] as well as neural activity [16]). However, these responses are less pronounced compared to when participants observed a human in pain. Specifically, functional magnetic resonance imaging (fMRI) data from the latter study provided initial evidence suggesting that the human brain responds differently to emotional displays by humans and robots via greater engagement within the right putamen (a brain region implicated in emotion processing) when participants observed videos depicting physical violence toward a person compared to videos depicting physical violence toward a robot [16].

Although these studies offer intriguing first insights into the emotional responses people experience when observing another person compared to a robot in apparent distress, an important consideration to bear in mind is that people have vast expertise observing and socialising with other humans but not robots, and these differences in exposure or familiarity could explain, at least in part, why emotional responses differ between humans and robots. As an estimated 1.5 million socially intelligent robots were sold between 2015-2018 (KPMG Advisory Report, 2016), this suggests that robots will become increasingly familiar in a variety of social settings, including schools, hotels, shops, and care homes [4]. To this end, it is vital to investigate if increased exposure to, or interaction or familiarity with social robots alters our emotional behaviour toward these machines, and to establish the functional changes at the behavioural and brain level [17]. This latter objective not only advances our understanding of human-robot relationships, but vitally, can also reveal fundamental and novel insights into the flexibility and adaptability of human social cognitive processes more generally [17,18].

In the present study, we examined how daily interaction with a socially-engaging entertainment robot across one week influences the extent to which people empathise with this robot, and how this compares to empathy directed toward another person. We used fMRI in conjunction with a robot socialising intervention to measure participants’ emotional (or empathic) responses while observing the human or robot’s response to either pleasant or unpleasant electrical stimulation, to examine not only empathy for pain, but also empathy for pleasure [16]. Identical fMRI sessions were run before and after participants were enrolled in a robot socialising intervention in which they took the robot home for five days to interact with it (**Figure 1A**). To map the flexibility of empathy for pain, we measured changes in repetition suppression of activity across the pain matrix network [8], comprising brain regions implicated in empathy for pain and emotion regulation, pre- and post-intervention. If empathising with a robot in pain or pleasure recruits similar neural mechanisms as when empathising with a human in pain or pleasure, viewing a human in pain followed by robot in pain should lead to reduced activity in the pain matrix. If dissimilar neural mechanisms are recruited for human and robot emotion perception, then we would not expect to see any suppression in activity (**Figure 1B**). We expected that socialising with a social robot should result in more overlap in neural mechanisms when observing a human in pain or pleasure followed by robot in pain or pleasure (or vice versa), leading to more pronounced repetition suppression of neural activity after socialising. Specially, the primary hypothesis we examined here is that repetition suppression for agents (robot and human), but not for emotions (pain and pleasure), should be more pronounced after compared to before the socialising intervention.

**Figure 1.**
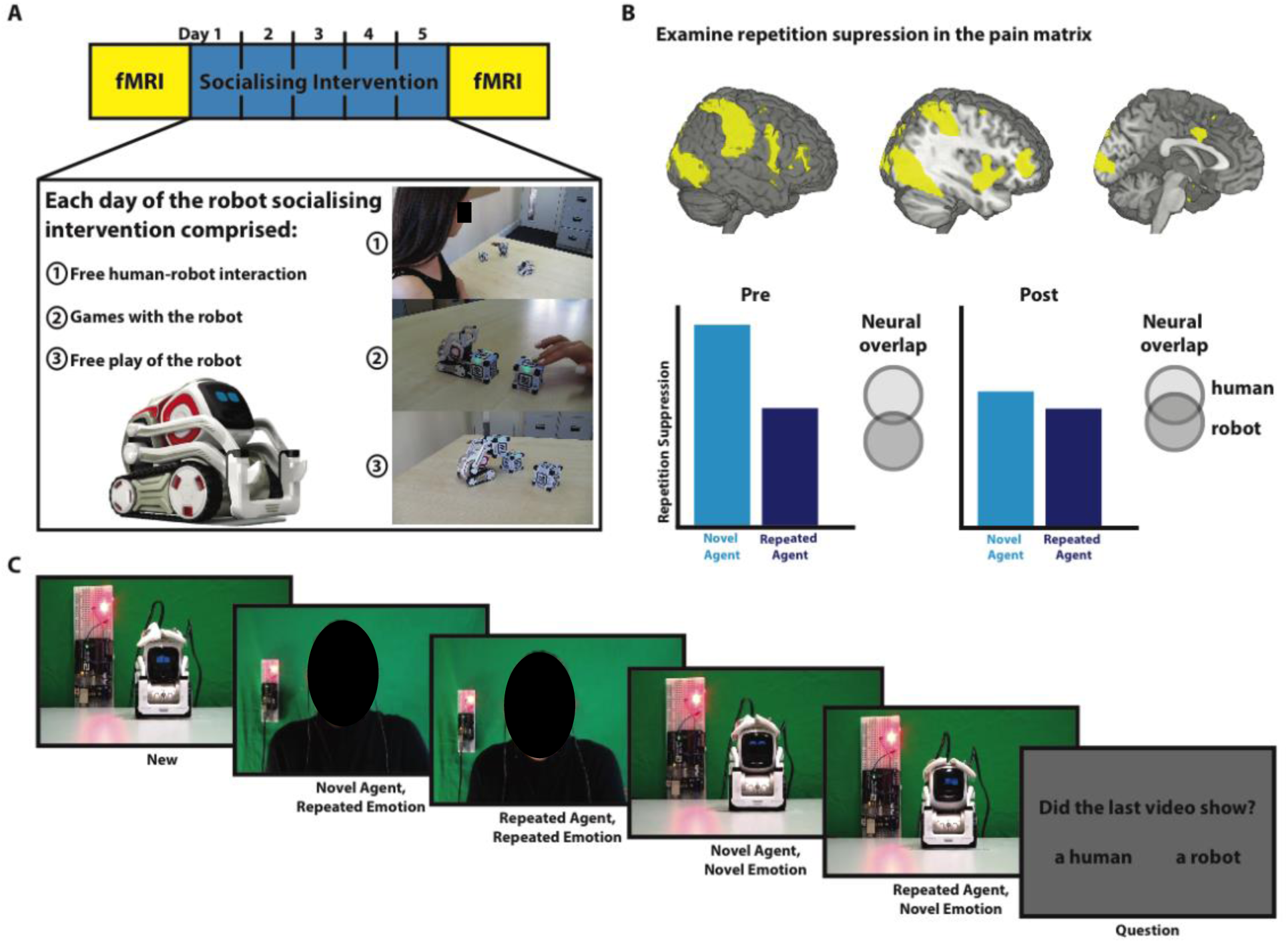
Overview of the study. **(A)** Schematic representation of the experimental design. Before and after the socialising intervention participant completed an identical fMRI session. After the first fMRI session, participants were enrolled in a five-day robot socialising intervention. Participants were given a small social robot and were instructed to interact at least 30 minutes each day. During the socialising intervention, participants were able to engage in free human-robot interaction (e.g., social play), play games (e.g. memory game), or observe the free play of the robot. **(B)** In the two fMRI sessions neural activation across regions of the pain matrix was measured using a repetition suppression design. The observation of a repeated agent, for example a human in pain followed by a human in pain, would suppress activity in the pain matrix. If empathy for a robot is dependent on similar or overlapping neural mechanisms as empathy for a human, the observation of a robot in pain followed by a human in pain would also lead to suppression of activity in these regions. If dissimilar or non-overlapping neural mechanisms are in place suppression of activity would be not be observed. Combining this repetition suppression paradigm with the socialising intervention allows to answer the question if socialising with a social robot would result in an increase overlap in neural mechanisms. **(C)** In the two fMRI sessions participants observed a human or robot in pain or pleasure. We employed a two factorial one-back repetition suppression design, with agent (novel or repeated) and emotion (novel or repeated) as factors. Each sequence started with a new video, followed by a novel or repeated agent showing a novel or repeated emotion. At the end of each sequence participants answered a question on the content of the last video (Did the last video show a human or robot? Did the last video show pain or pleasure?). Please note that images of the human agent have been obscured due to bioRxiv’s policy to not post manuscript that contain identifying material.

## 2. Materials and methods

### a) Preregistration and data statement

This study was pre-registered at www.AsPredicted.org, and the pre-registration can be found at http://aspredicted.org/blind.php?x=hb4cc7. We report all measures in the study, all manipulations, any data exclusions, and the sample size determination rule. All behavioural and functional region-of-interest data and stimuli are publicly available via the Open Science Framework, https://osf.io/9h4n7/?view_only=1a4a85b0d28942af8d5da2f308fc1d51, and the whole-brain group contrast maps can be found at NeuroVault [19], https://neurovault.org/collections/ZFRWTMDZ/.

### b) Participants

As determined before the start of the experiment, we collected data for twenty-eight participants. The sample consisted of 20 women and 8 men, aged between 18 and 27 years old, who were recruited from the Bangor University student population. All had normal or corrected to normal vision, and no history of neurological or psychiatric disorders. Participants were naïve to the goal of the experiment, were unfamiliar with the robot used in the study, and were not regularly exposed to robots in daily life, movies or TV shows (e.g. regular watchers of *Westworld, Humans*, etc.). Median engagement with robots in daily life on a scale from 1 (never) to 7 (daily) was 1 with an interquartile range (IQR) of 2, and median number of seen movies or TV shows that featured robots was 4 (2) out of 14 listed [20]. Participants received verbal and written information prior to the study, provided written informed consent before beginning any study procedures, and were reimbursed upon completion of the study. Final sample size was 26. One participant dropped out halfway through the study, while another participant was excluded due to excessive movement (spiking > 3mm) during fMRI. The final sample comprised 8 males and 18 females with a mean age ± SD of 19.85 ± 2.11 years. All study procedures were approved by the Bangor Imaging Unit and the Bangor University School of Psychology Research Ethics Committee (protocol number: 2017-16209) and carried out in accordance with the standards set by the Declaration of Helsinki.

### c) Design

Participants completed two fMRI sessions, pre- and post-interaction, which were identical with the exception that the functional localiser and anatomical scans were collected during one session only. In between the two scanning sessions, participants completed a robot socialising intervention wherein they were asked to interact with the robot daily for five days. The first fMRI session corresponded with the first day of the interaction, while the second fMRI session corresponded with the fifth and final day of the interaction (**Figure 1A**). Each fMRI session included four runs of trials that adhered to a 2×2 factorial design with novel or repeated agents and emotions (novel agent, novel emotion; repeated agent, repeated emotion; novel agent, repeated emotion; repeated agent, novel emotion; see **Figure 1C** and Stimuli and apparatus, below).

### d) Stimuli and apparatus

For the robot socialising intervention, we used a commercially available robot named Cozmo (**Figure 1A**; Anki, San Fransciso, CA, USA). This palm-sized robot (size: 5 × 7.25 × 10”) features a small head with a LED display (128 × 64 resolution), a VGA camera (30 FPS), four motors and >50 gears and can move its head (up/down), fork (up/down), and tracks (forward/backward and left/right). The human user is able to interact (e.g. free play) or play games (e.g. memory games) with the robot. The robot features an emotion/behaviour engine which, alongside external triggers, regulates its emotional reactions. Examples of external triggers are known or unknown individuals, pets, game events, or the edge of a surface or simply the lack of any external triggers. These external triggers affect the values in the emotion/behaviour engine, such as the frustration value or satisfaction value of the robot. The reactions of the robot are dependent on these values. For example, when the robot plays an animation or reacts based on a direct stimulus like winning a game, the intensity of the emotion displayed depends on the current values in the emotion/behaviour engine. Thus, if the frustration value is low, the robot will give a small negative reaction to losing a game, while if the frustration value is high the reaction will be more intense. These reactions affect the values in the emotion/behaviour engine in a similar dose-dependent manner. A low intensity reaction triggered by frustration brings down the corresponding factor only slightly, whereas a high intensity reaction will bring down the frustration value completely. When interacting with human the robot uses facial recognition to identify people and their basic facial expressions (for example, happy, sad, fearful and surprised expressions). It works together with an application on a compatible tablet device. We used the application version 1.5.0 that does not feature the extensive programming possibilities offered to the user by later versions (1.6.0 and up).

The stimuli used in the fMRI session consisted of eight 10s videos depicting a human or robotic agent appearing to receive painful or pleasant electrical stimulation to the head (**Figure 1C**; in reality, no stimulation was actually delivered to either agent). The human agent was a female actor and the robotic agent was the Cozmo robot used in the robot socialising intervention. To create the illusion that the agent was receiving either painful or pleasant stimulation, two wires connected to an electrical circuit were attached to the temples of the human agent and the head of the robotic agent. All videos started with the agent facing the observer with the two wires and electrical circuit visible. After ∼4 seconds, a red light-emitting diode (LED), positioned on an electrical circuit to the left of the agent’s head, lighted up and the agent exhibited a pain or pleasure response. This response lasts for approximately 4 seconds in length, after which the agent returns to a neutral facial expression. The intensities of the emotion ranged from mild pleasure or pain to extreme pleasure or pain. The emotional reactions displayed by the robot were triggered using the robot’s SDK (animations number: 27, 37, 146, and 215). For example, one pain reaction consisted of the robot rapidly rolling backwards, the LED ‘eyes’ tilting downwards, and raising its fork-lift style arms. One pleasure reaction involved the robot giggling, LED eyes tilting upwards, and the fork lift arms raising slightly, multiple times. The human actor was instructed to convey pain or pleasure and was made familiar with the range of movement and reactions of the robot. To portray pain the actor winced, clenched her teeth, raised her shoulders and tilted her head back. To convey pleasure, she smiled, widened her eyes, lowered her shoulders, and giggled. In order to increase believability of the pleasant stimulation, the actor was instructed to imagine it was like a gentle vibration on the temples that could be construed as pleasant. In order to create the illusion that this behaviour was in response to electrical stimulation, the actor was prompted by the experimenter to perform this behaviour as the LED was illuminated. The actor was then asked to reduce the intensity of the emotion and return to a neutral expression to reflect recovery from the stimulation.

To create a stimulus database that contained expressions of extreme and mild pain and pleasure expressed by the robot, three experimenters watched all animations within the robot’s repertoire (within the Cozmo application version: 1.5.0) and selected videos which appeared to convey pain or pleasure most clearly. The final set of videos was selected based on independent ratings of multiple videos across three online pilot experiments. First, the 11 videos were independently rated in an online pilot experiment. Participants (*n* = 19) rated each video on a continuum from ‘most painful’ (−5) to ‘most pleasant’ (+5). Based on the outcome of this pilot experiment, videos that were rated inconsistently were removed from the stimulus set. The remaining seven robot videos were rated by different participants (*n* =48) on the same continuum. In the final pilot experiment, four videos that corresponded to extreme and mild expressions of pain and pleasure as well as a neutral expression were selected from the stimulus set. Participants (*n* = 34) first watched and rated all robot videos, followed by watching and rating of videos of a male and female actor, each expressing a 7-step parametric shift from extreme pleasure to extreme pain.

Across all three pilot experiments, a parametric shift from extreme pain to extreme pleasure was observed (**Figure S1**). Results showed that the robot expressions in the four selected videos map onto the extreme and mild painful or pleasure categories: median rating (IQR) for extreme pain: −3.25 (2), mild pain: −1.5 (2.5), mild pleasure: 0.375 (3.5), and extreme pleasure: 2.55 (2.74) (**Figure S2)**. Ratings of the female actor showed less variability; therefore, these videos were selected for the main fMRI experiment: extreme pain: −4.75 (0.75), mild pain: −3.25 (1.19), mild pleasure: 3.13 (1.25), and extreme pleasure: 4.38 (1) (**Figure S3**). In the main experiment, participants only saw one robot and one human actor in all the videos.

### e) Robot socialising intervention

After the first fMRI session, each participant was issued a Cozmo robot and a Lenovo tablet (Lenovo Tab 4 or 7) with the Cozmo application preinstalled. Participants took the robot and the tablet home and were instructed to interact with the robot for at least 30 minutes per day across the next five days. No instruction was given to the type of interaction or play except for the instruction to invite a friend for a group interaction and play towards the middle of week. The application gives the user the possibility to unlock games (e.g. meet Cozmo game, quick tap game), features (e.g. pickup cube, fist bump), or complete daily goals (e.g. ‘win a game of quick tap with 100% accuracy’, ‘let Cozmo play uninterrupted for 10 minutes’). By completing daily goals, the human user earns energy, and earns bits and sparks rewards that can be used to unlock upgrades or features within the robot (games or features). Before each handout, the game history of the previous owner was erased. Each robot was therefore distributed to each participant in the same “naïve” state. During the robot socialising intervention, the application data were locked on the device to prevent sending the data to the developer. At the end of the robot socialising intervention, the data were extracted and stored for offline analysis. For each interaction a participant has with the robot, the application logs the state of the emotion/behaviour engine, the social behaviour of the robot, as well the amount, duration and type of the interactions between the robot and the human. In the present study, we used these log files to establish the quantity (number of games, duration of the interaction per day) and quality of the interaction (positive and negative emotions of the robot quantified by B.C., and ratio of self-initiation of the robot (requests made by the robot divided by total behaviours of the robot) between the human and the robot.

### f) Neuroimaging procedure

At the beginning of the first fMRI session, participants were introduced to the Cozmo robot. They were shown a brief videoclip in which a voiceover introduces Cozmo as a “robot with personality” followed by a brief overview of its key features and capabilities (e.g., face recognition, emotion/behaviour engine) by the voiceover, and the cofounder and president of Anki (0.00s – 0.56s of the video “Meet Cozmo, the AI robot with emotions”, https://www.youtube.com/watch?v=DHY5kpGTsDE). In the video, the robot can be seen reacting to a nearby hand, exploring its surroundings, expressing emotions via its LED eyes, and playing a game.

In each fMRI sessions, participants completed the same primary behavioural task, which adhered to a one-back repetition suppression design based on previous studies [21,22]. Participants observed sequences of five movies, composed of a first new movie followed by four videos that showed a novel or repeated agent with a novel or repeated emotion (**Figure 1C**). The order of the videos within the sequence was pseudo-randomized to create a balanced two-by-two factorial design (agent: novel and repeated, emotion: novel and repeated). The definition of each video, in terms of novel or repeated agent and emotion, is conditional on the previous video. Each video was part of all four conditions, counteracting any differences in low-level visual properties [22]. For instance, a video showing a robot experiencing pain can be a novel agent, novel emotion event in sequence 1, a novel agent, repeated emotion event in sequence 19, a repeated agent, novel emotion event in sequence 31, or a repeated agent, repeated emotion event in sequence 48. Each video was 10s in duration followed by a blank interval of 500ms. Participants were instructed to closely watch the videos and were told that they would see either a human or a robot receiving electrical stimulation that can be either painful or pleasant. To probe alertness, they answered a question (‘did the last video show a human or robot video?’ or ‘did the last video show pain or pleasure?’) at the end of the 10 out of 12 sequences per run. Participants responded with their left or right index finger and the type of question and answer positions were pseudo-randomized. The order of the sequences was randomized, and each sequence was followed by a fixation cross (two, three, or four seconds in duration; pseudo-randomized). Each run consisted of 12 sequences in total with a total duration of 12.1 min. Participants completed four runs with 192 trials in total (48 trials per condition).

We used a functional localiser to identify regions of the pain matrix [23]. At the start of either the first or second scanning session (depending on scanning schedule timing), participants passively viewed a short 5.6 min animated film (‘Partly Cloudy; https://www.pixar.com/partly-cloudy#partly-cloudy-1). This movie includes scenes depicting pain (e.g., an alligator biting the main character) and events that trigger mentalizing (e.g., the main character revealing its intention). Stimuli were presented using Psychopy 3.0.11 [24] (functional task) and Octave 4.0.0 (http://www.octave.org) with Psychophysics Toolbox 3.0.11 [25-27] (functional localiser), and projected onto a MR safe BOLD screen (Cambridge Research Systems: http://www.crsltd.com/), visible to participants via a mirror mounted on the head coil.

After each fMRI session, participants rated each video seen during scanning four times on scale from −5 (pain) to +5 (pleasure), and were asked to rank-order the robot videos as well as the human videos from most painful to most pleasure. In addition, they completed questionnaires to measure the shift in attitudes to and perception of robots (Negative Attitudes towards Robots Scale (NARS); [28,29], the Godspeed questionnaires; [30], and the agency in robots questionnaire; [31], anthropomorphism [32,33]), and self-other overlap with the robot (Inclusion of the Other in the Self; [34]). At the start of the first fMRI session, participants completed questionnaires to probe empathy (Interpersonal Reactivity Index (IRI); [35], and daily exposure to robots [20]). All questionnaires, except for the Inclusion of the Other in the Self (IOS) and the daily exposure to robots, were collected for a different research project and are thus not considered further in this paper. In the IOS, participants rated their perceived self-other overlap between them and four other agents (the robot, person in the video, a close friend, and a stranger) on a scale from 1 (no overlap) to 7 (complete overlap).

### g) fMRI data acquisition

Data were acquired with a 3-Tesla Philips Achieva full-body MRI scanner using a SENSE phased-array 32-channel head coil. Participants were provided with earplugs and headphones to attenuate scanner noise and foam padding was used to reduce head movements. Functional images were acquired using a T2*-weighed single-shot echo planar image sequence [functional task parameters: repetition time (TR) = 2500 ms, echo time (TE) = 30 ms, number of slices per volume = 36, 3 mm in-plane isotropic resolution, no gap, flip angle (FA) = 90°, field of view = 213 × 213 mm^2^, matrix size: 240 × 224, ascending slice acquisition, anterior-posterior phase encoding, number of volumes per run 292; functional localiser: TR = 2000 ms, TE = 30 ms, number of slices per volume = 32, 3 × 3 × 3.5 voxels, no gap, FA = 83°, field of view = 240 × 240 mm^2^, matrix size: 240 × 224, ascending slice acquisition, anterior-posterior phase encoding, number of volumes per run 180]. For the first three sessions, the response screen was not fixed resulting in slightly different numbers of volumes per run (which were taken into account during preprocessing). At the end of the first or second session (depending on timing and scanner schedule), high-resolution structural images for each participant were collected using a three-dimensional T1-weighted imaging sequence scan [parameters: 1 mm isotropic resolution, TR = 12 ms, TE = 3.5 ms, FA = 8°, field of view = 240 × 240 mm^2^].

### h) fMRI preprocessing and analyses

Image preprocessing and analyses were carried out using SPM12 (Wellcome Trust Centre for Neuroimaging, London; [36]) in MATLAB R2017a and 2017b (Mathworks, Natick, MA, USA). Preprocessing of functional data per session consisted of realignment and unwarping, slice time correction before realignment, and spatial smoothing with a 6 mm kernel (5 mm for the functional localiser; [23,37]). Images were normalized to the Montreal Neurological Institute (MNI) template with a resolution of 3 mm.

Three design matrixes were created for the main repetition suppression analysis, non-repetition suppression analysis, and the functional localiser. Motion predictors were included as predictors of no interest in all three design matrixes. The onset and duration of each event was specified and convolved with the standard hemodynamic response function. The design matrix for the one-back repetition suppression design contained seven task regressors, four for each condition, and regressors for the new movie, question, and rest period in between the sequences. Besides the four repetition events, the main first-level contrasts calculated were a) main effect of agent (novel > repeated), and b) main effect of emotion (novel > repeated). For the non-RS analyses the regressors were recoded into discrete events, cf. a robot or human experiencing pain or pleasure. This resulted in four regressors for the videos, human or robot experiencing pain or pleasure, and the two additional regressions (question and rest period). Main first-level contrasts calculated were a) main effect of agent (human > robot), b) main effect of emotion (pain > pleasure), and c) interaction between agent and emotion (human (pain > pleasure) > robot (pain > pleasure).

To measure the pre-post change in repetition suppression, we used group-constrained subject-specific analyses [38,39]via the spm_ss toolbox (http://web.mit.edu/evelina9/www/funcloc.html) implemented in SPM. This approach increases sensitivity and functional resolution and decreases ambiguity in the selection of functional regions of interests (fROIs) [40]. Subject-specific functional ROIs for the pain matrix were created by masking the contrast images from the functional localiser per participant (*p* < .001, k = 5) with independent parcels. Independent parcels for 7 replicated brain regions in the pain matrix were used from Richardson et al. [37]. These parcels consisted of the anterior middle cingulate cortex, and bilateral second somatosensory cortex, insula, and middle frontal gyrus (**Table S1)**. We selected the top 10% of the activated voxels in the localiser contrast to create subject-specific fROIs constant in size across participants [41]. These subject-specific fROIs were used to extract the signal for the functional task. Upon extraction, we calculated the pre-post change per repetition suppression condition for each of the seven identified regions of the pain matrix. For example, the blood oxygen level-dependent response for the repeated agent, novel emotion condition pre-interaction minus suppression of the blood oxygen level-dependent response for the repeated agent, novel emotion condition post-interaction. A repeated measures analysis of variance (ANOVA) with agent (2) and emotion (2) as within-factors was used to assess the socialising induced change in repetition suppression. Positive values indicate more repetition suppression post-socialising, whereas negative values indicate less repetition suppression post-socialising. A similar analysis was performed for the discrete coded events approach, where positive values indicate less activation post-socialising and negative values indicate more activation post-socialising.

Besides group-constrained subject-specific analyses, we also ran whole-brain random-effects analyses for the repetition suppression design, focusing on main effects of agent and emotion, and for the design featuring discrete coded events, focusing on main effects of agent (human > robot and robot > human), as well as agent by scan session interactions. The initial single voxel threshold for these maps was set to p < 0.001, k = 10 voxels. We focus on clusters that meet the cluster-level false-positive rate of 5% (p*FWE-corrected* < 0.05), and visualise activations from the whole-brain repetition suppression analyses at the uncorrected threshold for further clarity (due to the subtle nature of these findings). Anatomical localisation of all activations was assigned based on consultation of the Anatomy Toolbox in SPM [42,43].

## 3. Results

### a) Robot socialising intervention

Across the five days of the socialising intervention, 17 of the 26 participants interacted more than the combined duration of the requested duration per day (150 min). Three participants interacted between 120 and 150 minutes and six participants did not meet the preregistered cut-off of ≥2 hours total interaction time. Mean duration ± sd interacted with the robot was 191.8 ± 86.8 min, acquired over 6.5 ± 1.9 sessions with a duration of 29 ± 8.70 minutes (**Figure 2A**). Participants completed 17.42 ± 9.20 games with the robot and ratio of self-initiation of the robot was 0.007 ± 0.006. Importantly, the interpretation of the behaviour of the robot is dependent on the human user, with the behaviour and expressions of the robot consisted predominantly of mixed emotional signals, 88.6 ± 2.12 %, with few clear positive, 5.11 ± 1.72, and negative signals, 6.32 ± 1.59. As the number of participants failing our preregistered cut-off surpassed our expectations, we ran the main fMRI analyses with and without these participants to determine the extent to which limited interaction time with the robot influenced neural engagement during the fMRI task. Exclusion of these participants did not influence the outcome of the fMRI analyses.

**Figure 2.**
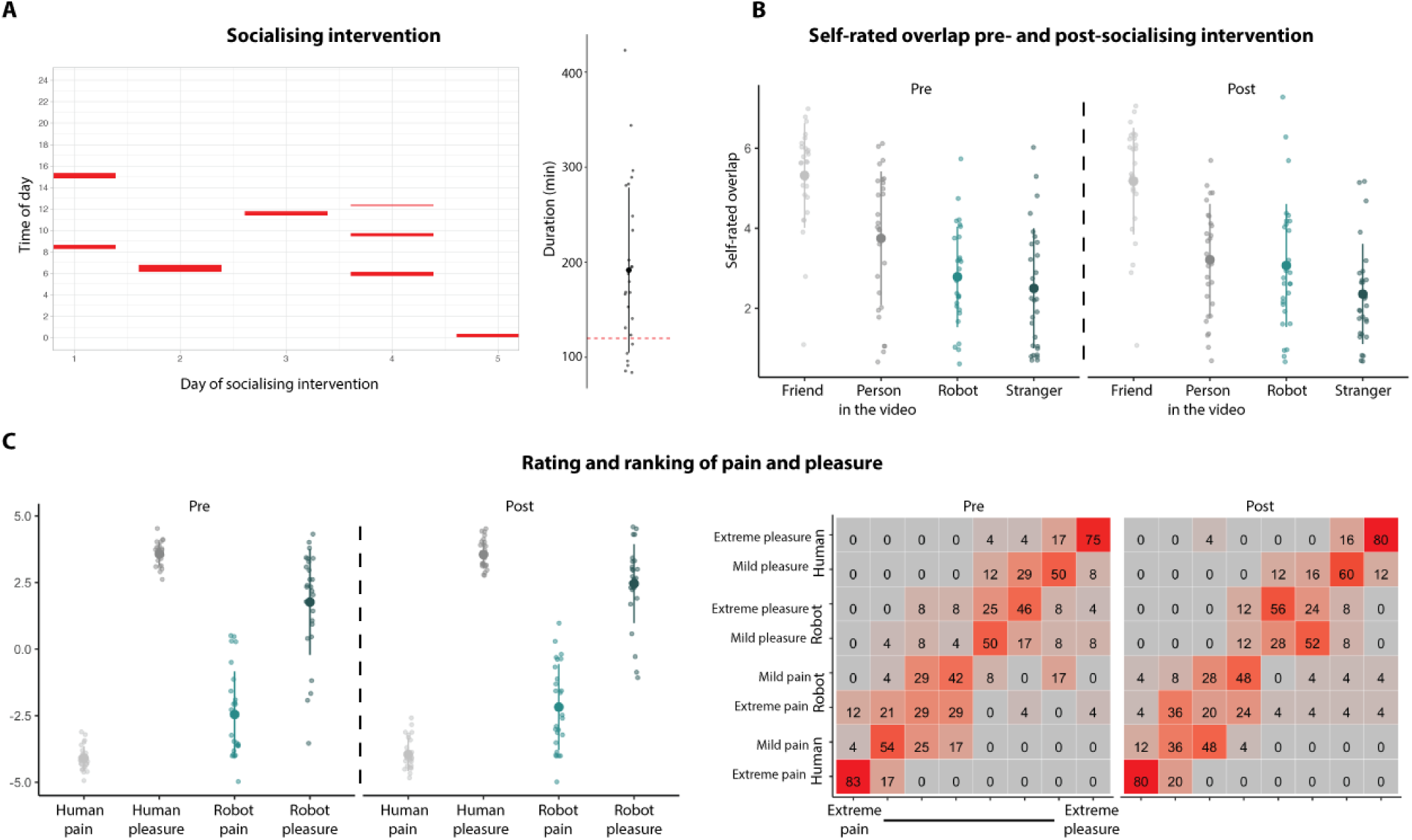
Socialising intervention and behavioural measures. **(A)** Participants were instructed to interact at least 30 minutes each day across the five-day socialising period. Session and duration for one participant in shown. Red lines denote the duration of each session. Most participants interacted with the robot longer than the required duration (150 minutes) or the cut-off (120 minutes, red dashed line). **(B)** Self-rated overlap with a friend, stranger, the robot or with the person in the video pre and post-socialising intervention. The socialising intervention did not influence self-reported overlap with the robot. **(C)** Rating and ranking of the human and robot experiencing pain and pleasure pre- and post-socialising. All videos mapped onto the expected pain and pleasure categories. Across the two sessions, the pain and pleasure experienced by the robot was rated as less intense. Mean and standard deviation and individual data is shown, and percentage of the video rated per category is shown for the ranking. Note, the analysis of the rating data is performed on absolute values.

Self-rated overlap with the robot did not increase after the socialising intervention, median (IQR) pre-socialising: 3 (1.75), post-socialising: 3 (2) (**Figure 2B**). A significant main effect of agent, *F*(3, 75) = 45.62, *p* < .001 η^2^ = 0.65 was observed, with self-rated overlap with a friend rated higher compared to other agents. No main effect of session, *F*(1,25) = 0.47, *p* = .50, η^2^ = 0.02, or interaction between session and agent was observed, *F*(3, 75) = 2.63, *p* = .06, η^2^ = 0.10.

### b) Subjective rating of pain and pleasure

Ratings and rank-order of videos mapped on to the distinct categories (**Figure 2C**). To carefully consider the effect of socializing on the rating of both pain and pleasure, we took the absolute value of the ratings. A repeated measures ANOVA with agent × emotion × session, revealed a main effect of agent, *F*(1,25) = 64.01, *p* < .001, η^2^ = 0.72. Robotic expressions were rated as less intense, with a median rating of 2.6 (SD = 1.9) compared to human expressions, which had a median rating of 3.9 (SD = 0.9). While an emotion by session interaction was observed, *F*(1,25) = 4.33, *p* = .05, η^2^ = 0.15, suggesting that pre-socialising pain ratings were higher compared to pre-socialising pleasure ratings, post-hoc comparisons did not survive correction for multiple comparisons.

### c) Task performance

No participant failed the preregistered attention check (<32 responses out of 40 questions per session). Importantly, performance did not differ between sessions. Accuracy for questions probing the agent appearing in the last video was higher, 0.99 ± 0.03 than for questions probing the emotional content of the last video, 0.87 ± 0.10, *F*(1, 25) = 77.42, *p* <.001, η^2^ = 0.76.

### d) Group-constrained subject-specific fROI analysis

We successfully identified the functional region of interest in all participants (**Figure 3A**). A repeated measures ANOVA with agent (repeated or novel) and emotion (repeated or novel) was performed on the pre – post-socialising difference score to test the preregistered hypothesis that repetition suppression for agents, but not for emotions, would increase after the socialising intervention. In contrast to our predictions, no main effect for agent was observed (**Figure 3B**). Repetition suppression of activity in the pain matrix when observing a human in pain followed by a robot in pain or vice versa did not increase after the socialising intervention. Moreover, no significant main effect of emotion or an agent by emotion interaction were observed in the pain matrix (**Table S2** and **S3**).

**Figure 3.**
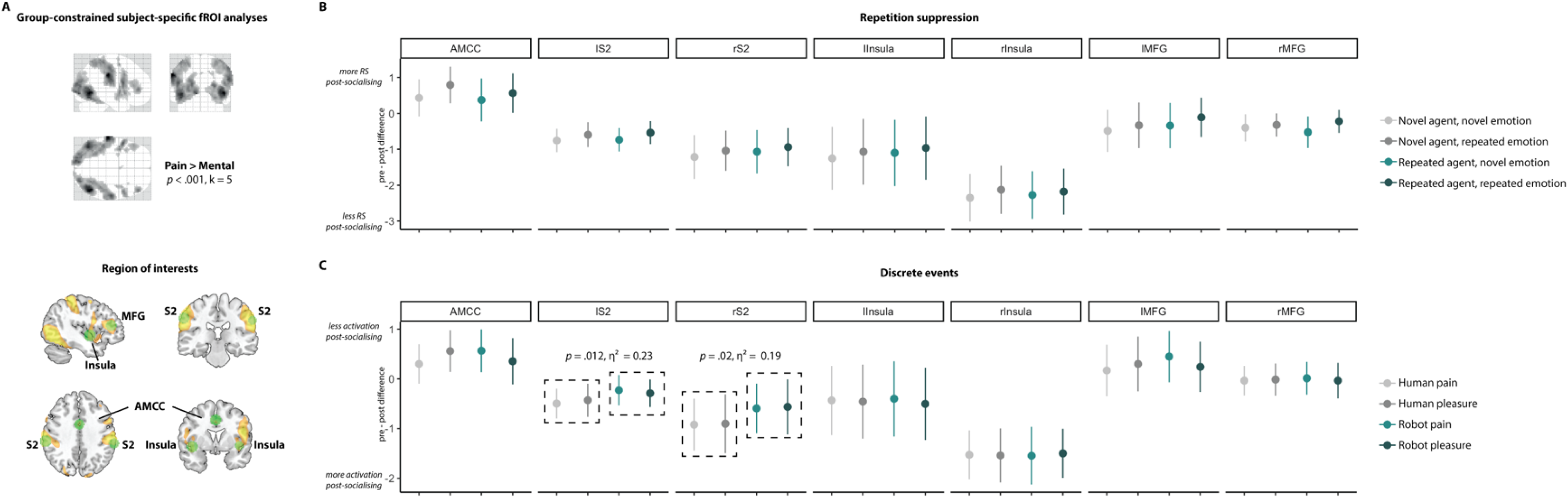
Group-constrained subject-specific functional region-of-interest approach and results. **(A)** We identified seven core regions of the pain matrix by combining an independent localiser with parcels derived from a previous study. Top figure shows group map for the pain vs. mental contrast for the functional localiser in the study sample (*p* < .001, extended cluster threshold of 5). Bottom figure shows the same group contrast maps with the parcels overlaid. Seven regions were identified for each participant based on the interjection of the parcels with their individual contrast map for the functional localiser. **(B)** Results for the repetition suppression analysis. No main effects for agent, emotion or interaction between agent and emotion were observed. **(C)** Results for the discrete event-related analysis. A main effect of agent was observed in the bilateral S2, suggesting a smaller increase in activity post-socialising when observing a robot experiencing pain or pleasure compared to when observing a human experiencing pain or pleasure. Mean and standard error of the mean is shown.

To map the possible emotion-dependent effects of socialising on activity in these regions we used a similar fROI approach but with the one-back repetition suppression design recoded into discrete, emotional events (e.g. a robot or human experiencing pain and pleasure). This event-related analysis revealed a main effect of agent in the bilateral secondary somatosensory cortex (S2), left S2: *F*(1, 25) = 7.41, *p* = .01, η^2^ = 0.23, right S2: *F*(1, 25) = 6.01, *p* = .02, η^2^ = 0.19 (**Figure 3C**). The increase in activity post-socialising was lower when observing a robot experiencing pain or pleasure compared to when observing a human experiencing pain or pleasure. No other effects were observed (**Table S4** and **S5**).

### e) Whole-brain analysis

As stated in our preregistration, we planned secondary random-effects analyses at the whole-brain level to evaluate a main effect of agent (novel > repeated), and a main effect of emotion (novel > repeated) with the repetition suppression design, for both the pre- and post-socialising scan sessions. We further explored whole brain results with the discrete coded event-related design, which enabled us to evaluate how observing human pain and pleasure compares to observing robot pain and pleasure (and vice versa), as well as any interactions between scanning session and how the different agents are perceived. For the RS design, we found several brain regions to be sensitive to novel agents (main effect of agent: novel > repeated agent, holding emotion constant), in both the pre and post-socialising scan sessions. Specifically, before the socialising intervention, bilateral precuneus and left early visual areas responded robustly to novel agents (**Figure 4A** and **Table S6**). After the socialising intervention, no cluster-corrected regions showed a sensitivity to novel agents, but bilateral medial temporal regions showed more subtle sensitivity to novel agents (**Figure 4A** and **Table S6**). The whole brain analysis evaluating the main effect of emotion revealed no cluster corrected brain regions sensitive to novel emotion displays in either the pre- or post-socialising scans, but did reveal subtle modulation of the middle temporal gyrus in the pre-socialising scan only (**Figure 4B** and **Table S6**).

**Figure 4.**
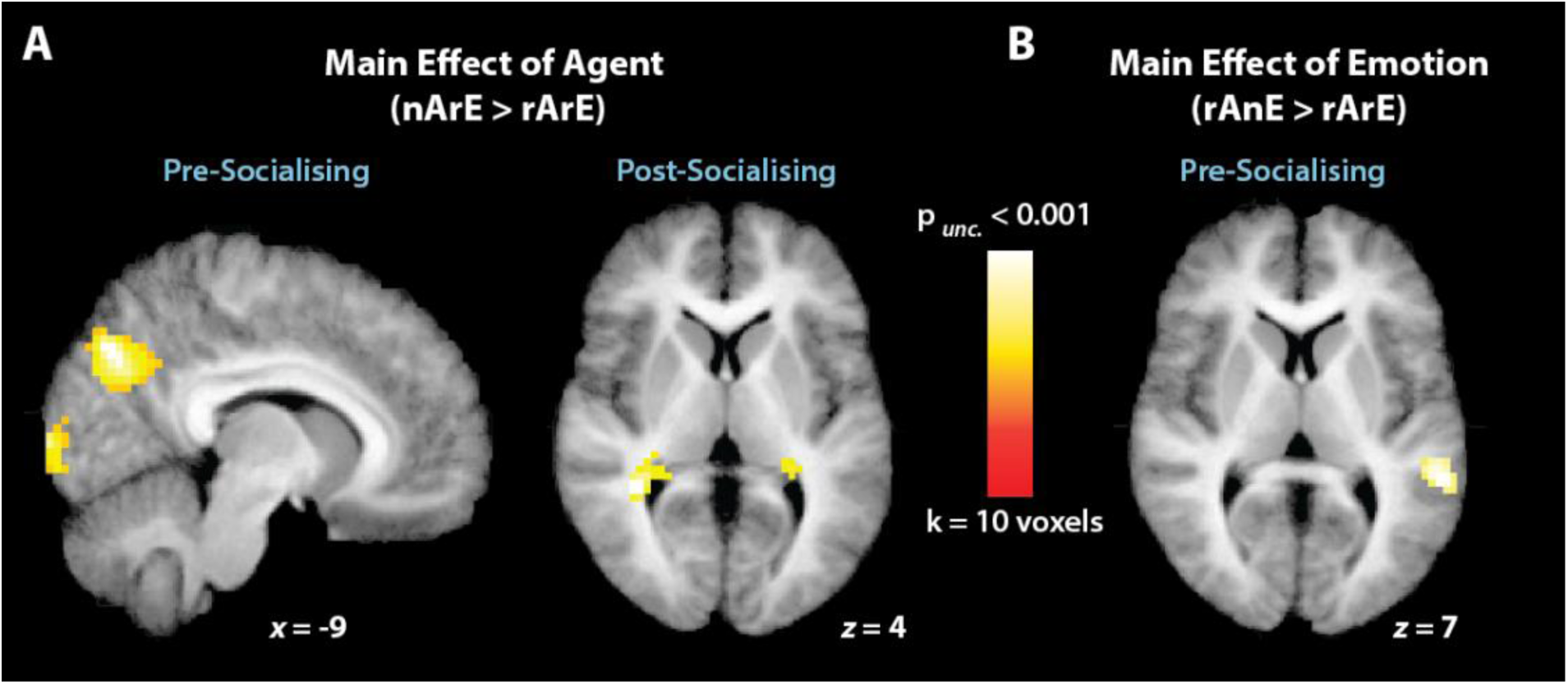
Whole brain analyses of main effects from repetition suppression design, broken down by session (pre- and post-socialising). **(A)** Illustration of brain regions sensitive to the appearance of a novel agent, while the emotion content of observed videos was held constant. Pre-socialising, this included robust (cluster-corrected) engagement of the precuneus and superior occipital gyrus, while post training, more subtle (non-cluster corrected) activations emerged within the medial temporal lobes. **(B)** Illustration of the main effect of emotion, while agent was held constant. This contrast yielded subtle (non-cluster corrected) engagement of the middle temporal gyrus bordering on the inferior parietal lobule.

Next, we evaluated whole brain analyses using the discrete-coded event-related factorial design. As we were most interested in how perception of human and robotic agents compares (more so than comparisons between positive and negative emotion displays), these analyses exclusively focused on the main effect of agent. As illustrated in **Figure 5A**, both before and after socialising, observing a human agent compared to a robot agent led to more robust engagement of bilateral medial posterior occipital cortices, including the precuneus and calcarine cortex. The inverse contrast revealed broad activation of occipitotemporal, parietal and premotor cortices, across both pre- and post-socialising scan sessions (**Figure 5B** and **Table S7**). Finally, all interactions between agent type and scan session were evaluated. The interaction examining brain regions more responsive to watching a robot compared to a human during the pre-socialising scan compared to the post-socialising scan was the only interaction to yield robust results. Specifically, this interaction showed a large cluster within the left superior parietal lobule, bridging the intraparietal sulcus, to respond more strongly to robots than humans during the pre-socialising compared to the post-socialising scan session (**Figure 5C** and **Table S7**).

**Figure 5.**
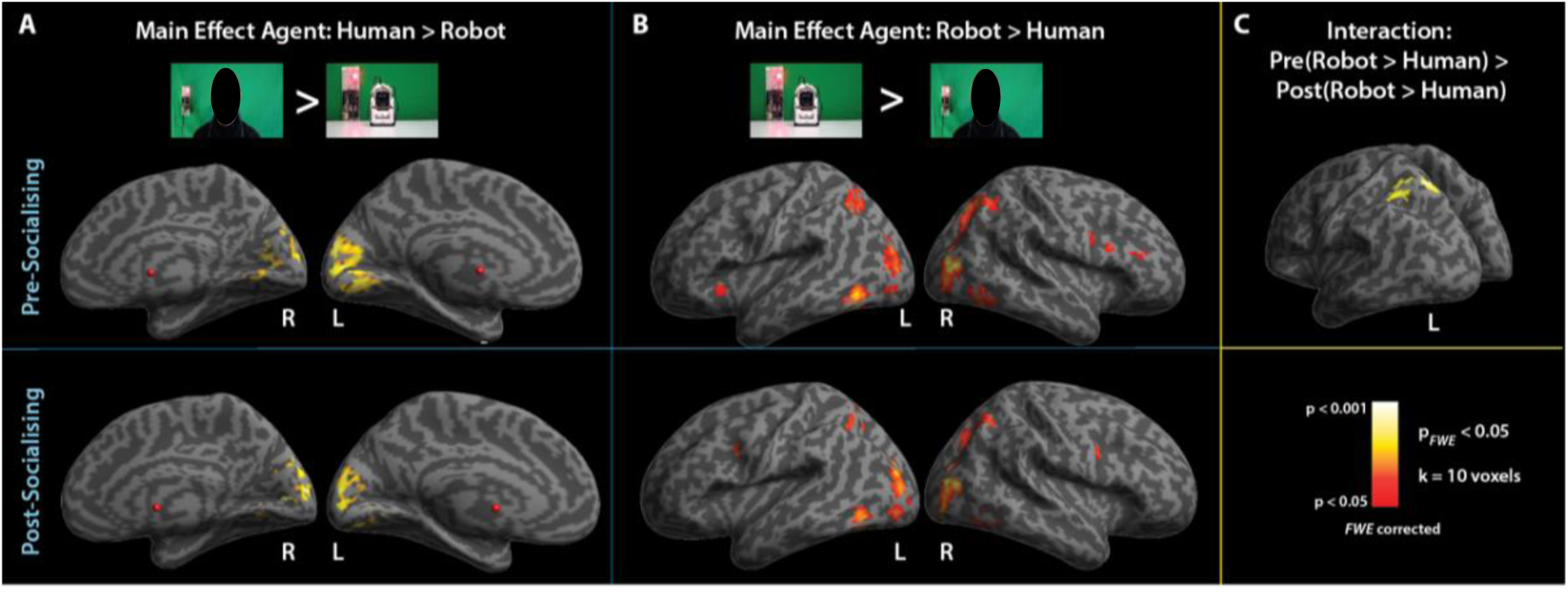
Comparison of the observation of human pain and pleasure and robot pain and pleasure. **(A)** Illustration of the main effect of watching videos featuring a human actor compared to a robot, during both pre- and post-socialising fMRI sessions. Early visual regions on the medial surface of the posterior occipital lobe emerged from these contrasts during both scanning sessions. **(B)** Main effect of watching videos featuring a robotic agent compared to a human actor, across both pre- and post-socialising sessions. Bilateral engagement of occipitotemporal, superior parietal and ventral premotor cortices/inferior frontal gyrus emerged during both of these contrasts. **(C)** One region within the superior parietal lobule, spanning the intraparietal sulcus, showed a more robust response to the robot compared to the human during the pre-socialising scan session compared to the post-socialising scan session. Please note that images of the human agent have been obscured due to bioRxiv’s policy to not post manuscript that contain identifying material.

## 4. Discussion

In the present study, we sought to investigate whether interacting with a robot across one week impacts empathy towards this robot at both behavioural and brain levels. The overall expectation was that a robot socialising intervention would result in an increase in overlap in mechanisms supporting empathy for humans and robots. Using both subjective and objective measures, our findings failed to support this hypothesis.

### a. Socialising with an entertainment robot does not markedly increase social feelings toward the robot

While the majority of the participants successfully completed the socialising intervention, no increase in self-rated overlap with the robot was observed. Participants reported similar levels of perceived overlap with the robot as with the human agent or a stranger, but this was not influenced by the socialising intervention. Similarly, ratings of observed pain and pleasure in the robot did not increase after the socialising intervention. These findings might lead us to ask to what extent socialising interventions, such as the one employed here, can actually influence social cognition. While research on repeated social interactions with artificial agents such as social robots is still limited, two early studies suggest that repeated exposure or interaction can indeed influence social cognitive processes [44,45]. Press and colleagues [44] tested the impact of repeated exposure to robotic actions on automatic imitation (a phenomenon that describes the influence of observed actions on executed actions) of robotic actions. Before repeated exposure, participants showed a human bias in automatic imitation, that is, the influence of observed human actions was greater than the influence of observed robotic movements. However, after a training session in which participants repeatedly made simple hand movements while observing similar robotic actions, this human bias disappeared. A different study by Tanaka and colleagues [45] showed that repeated interaction with or exposure to a robot can influence social cognition beyond sensorimotor processes. In this study, a small robot was deployed in a classroom of 18- to 25-month old toddlers for 5 months. Not only did the quality of interaction increase over the intervention, but toddlers also displayed similar behaviours toward the robot as to their peers (e.g., touch, care-taking and play). While these results are promising, no direct measures of individual social cognition were reported (and naturally, a five-month socialisation intervention with 18-25 month olds is quite different from the one-week socialisation intervention with young adults used here).

The study by Press and colleagues [44] already hinted at a common denominator in successful training or intervention studies, the focus on one well-defined aspect of social cognition. For instance, previous training studies successfully probed the flexibility of sensorimotor cortical engagement with training of dance sequences or other skilled actions [46,47]. In the present study, we focussed on real social interactions with an embodied robotic agent [17,18,48], and the training intervention took place in participants home, not in a supervised and tightly controlled laboratory environment. While this increases the ecological validity and engagement of participants, a downside is that it also increases inter-individual variability. As we did not constrain the behaviour of the robot, the interaction scenarios could vary wildly between participants. For example, one person might have predominantly played memory games with the robot, while another participant enjoyed more free play by the robot. Future studies should keep these limitations and benefits in mind, and perhaps make use of combination between a data-driven approach to distil common features of human-robot interaction and interventions of distinct aspects of social cognition.

### b. Establishing a role for the pain network in empathy toward robots, and the extent to which this changes with experience

Contrary to our main neuroimaging hypotheses, we found no increase in repetition suppression of activity in brain regions implicated in empathic responding to distress in others. Similarly, we failed to find evidence of emotion-dependent impact on activity within these brain regions. Overall, these results suggest that a one-week socialisation intervention with a robot does not lead to an increase in neural overlap for human and robot distress. One small effect we found was a smaller increase in bilateral S2 activity post-socialising for robotic pain and pleasure compared to human pain and pleasure. While replication of this finding will be necessary to establish its reliability, these effects converge with findings from other studies on the flexibility of pain matrix activity in response to another individual’s distress. Activation in parts of the pain matrix is modulated by expertise [49], social context [50]and group membership [50,51]. For instance, activation in this network was decreased for member of outgroups (e.g. rival football fans) [50,51]. Can activity in these regions also be upregulated? One study provided for first evidence for this by showing that a short empathy training resulted in increased activity in core regions of the pain matrix, the AMCC and insula, during the observation of suffering [52]. While these findings attest to the flexible nature of brain regions supporting empathy, the question remains if and how interactions with artificial agents shape engagement of these regions.

One caveat that should be mentioned is the ostensible functional role ascribed to the collection of brain regions that form the pain matrix. While several empirical and theoretical accounts suggest a role of the pain matrix in empathy [8,53,54], several others challenge this notion [55-57]. A contrasting account suggests that activity in parts of this network support the detection of salient events in the environment, and behavioural responses to these events [55]. For instance, regions that correspond to the pain matrix, including the anterior cingulate cortex, second somatosensory cortex, and insula, are activated in response to a broad range of salient events, regardless of nociceptive qualities, thus calling into question the selective nature of the pain matrix. While we used an established functional localiser [23,37], at this stage, we believe it best to refrain from making conclusive statements on the functional nature of the activation in this network, and await further investigations [53,57].

While people readily ascribe human-like characteristic to artificial agents like the one used in the present study, an intriguing question remains as to whether empathy can be felt toward artificial agents. Empathy is a multifaceted concept that consists of a variety of phenomena that differ in complexity ranging from mimicry to perspective taking [5]. Results from both qualitative and quantitative studies suggest that people can indeed to some extent feel empathic towards robots [15,58,59]. While we likely tapped into an affective or vicarious component of empathy in the present study, feeling empathy toward a robot might assume the presence of emotions in that robot (e.g. pain, suffering). A previous study found that people deem a robot capable of phenomena related to agency (e.g., memory, planning), but not experience (e.g., rage, hunger) [60]. However, people also accurately identify emotions expressed by a robot or other artificial agent, and neural networks are similarly engaged when perceiving emotions expressed by humans and robots [14,17]. An intriguing question that arises from this prior work concerns the extent to which explicit knowledge on the limitations of artificial agents’ capacity for experience-related phenomena, such as pain or suffering, impact bottom-up processes, like empathy for pain. We urge future studies to explore these questions further, perhaps using different measures of social cognition, to further elaborate the impact of socialising with a robot on the scope and limitations of the attribution of socialness to these agents.

### c. Insights gained from whole-brain analyses of agent and emotion perception

In addition to our main contrasts of interest within the functionally localised pain matrix, we performed several exploratory whole brain analyses to evaluate how perceiving dynamic displays of human- and robot-expressed emotion engages social perception more broadly. The main effect analyses of the repetition suppression design revealed that observing a novel agent resulted in engagement of posterior parietal and early visual regions before the socialising intervention, which shifted to more middle temporal/parahippocampal cortices during the post-socialising scan. While the findings from the first scanning session are broadly consistent with previous work examining how the brain codes novel identities [61], the finding that novel agents engage more parahippocampal cortex after a socialising intervention raises potentially interesting questions about whether particular memory processes might be engaged after a brief socialisation intervention. The contrast examining the main effect of emotion within the RS design revealed a cluster spanning the middle temporal and inferior parietal cortices to respond more robustly to novel emotion, which might reflect additional engagement of attentional resources based on the shifting of emotional valence between videos [62,63]. The whole brain analyses of the discrete-coded event-related design revealed additional insights into how participants perceive human compared to robotic agents and actions. In general, the finding of stronger engagement of early visual regions when observing human videos and more robust engagement of frontal, parietal, and occipital temporal cortices when observing robot videos complements and extends previous work comparing how we perceive human and robotic actions [64,65].

### d. General limitations and ideas for future research

While our data failed to support our primary hypothesis, a number of considerations are worth bearing in mind that might help improve future research efforts along these lines. By mapping changes across two identical fMRI sessions before and after the socialising intervention with a trained (robot) and untrained (human) agents, the current study followed a similar procedure as previous training studies on sensorimotor learning and action observation (e.g., [46,66]). While using a pre- and post-intervention approach already limits the influence of potential confounding factors that might arise if a single scan session is performed post-intervention only, a question could nonetheless arise as to whether the repetition of the task or time between the two fMRI sessions could impact the present results. Future studies may wish to consider including a control group without a robot socialising intervention or a variable washout period between the two fMRI sessions to counteract these potential confounds. One intriguing solution to circumvent some of these issues has been employed by Tanaka and colleagues [45] and involves the manipulation of the robot’s behaviour during the socialising intervention. After the first 27 sessions where the robot had its full behavioural repertoire, the robot was re-programmed to perform repeated and restricted behavioural sequences with limited contingency. This would allow to tap into the complexities of the long-term interactions and the factors that influence bonding and changes in social cognition, while at the same time restrict the influence of potential confounding factors on the experimental results. One final issue that is not specific to the present study, but holds for the literature at large, is the possibility of agent-dependent effects and confounds. As every robot brings different stimulus cues (such as what they look like and how they move) to the interaction, it might be possible that individual studies tap into idiosyncratic effects tied to the specific robot used for each study. As such, questions remain as to whether the current findings, and the socialising or long-term interaction effects reported in the literature more broadly, generalize across robotic agents. Future studies may wish to include multiple robotic agents to study the specificity and generalizability of the impact of socialising or long-term interactions with a robot on human social cognition.

While our sample size is by no means on the small side of many published fMRI studies, this does not change the fact that most neuroimaging studies are statistically underpowered, consequently reducing the chances of detecting true effects [67,68]. On top of this, studies that examine effects of training or other types of interventions that take place across time are often searching for subtle effects that are even more elusive than what is being examined during single-session fMRI studies [69]. While we absolutely acknowledge that a larger sample size would have been ideal to test the current hypotheses, we also argue that time-intensive training invention studies, such as the one described here, are valuable for understanding how cognitive and perceptual processes change across time, and are worth considering even if there are power considerations [70]. Future studies seeking to examine how socialising with artificial agents shapes social perception and cognitive processes may wish to deploy longer or more intensive socialising or training procedures, as well as experiment with alternative methods for examining the neurocognitive mechanisms supporting social engagement with human and robotic agents. As an example, multi-voxel pattern analysis can provide a more sensitive window into the flexibility of social cognition while interfacing with artificial agents. This approach has been successful in mapping subtle changes in neural representation within the frontoparietal cortex after a training intervention to foster observational learning [71], and within the auditory cortex and lateral frontal cortex after training to discriminate communicative calls of monkeys [72]. The exciting combination of real interactions with embodied agents and advanced neuroimaging analyses provides promising new avenues to study the functional changes in engagement of brain regions at the core of everyday social interaction.

## 5. Conclusion

In sum, a better understanding of the mechanisms and consequences of how humans establish social relationships with robotic agents will be vital if these machines are to take on increasingly social roles within human society. While the present study did not find evidence that a one-week socialisation intervention with an engaging social robot led to a more human-like empathic response to seeing this robot in pain, it is vital that future research examines longer term and richer socialisation interventions to determine with more certainty whether the human brain might ultimately come to perceive and relate to robots in a similar way as humans.

## Supporting information

## Authors’ contribution

ESC and RH with KAR conceived the study. ESC and RH with KAR, JP, and ST designed the study. ESC, KAR, JP and ST performed the experiments. BC provided vital robotics programming and technical support and expertise throughout the project. ESC and RH analysed the data and wrote the paper.

## Acknowledgements

The authors would like to thank Kohinoor Darda for assistance during fMRI analyses, Hilary Richardson for providing the derivatives for the group-constrained subject-specific functional regions of interest analyses, and the Social Robots team for engaging in helpful discussions at an early stage of the project (particularly Laura Jastrzab Binney) and for providing critical feedback on the manuscript.

