## Supplementary material for "A neurocognitive investigation of the impact of socialising with a robot on empathy for pain"

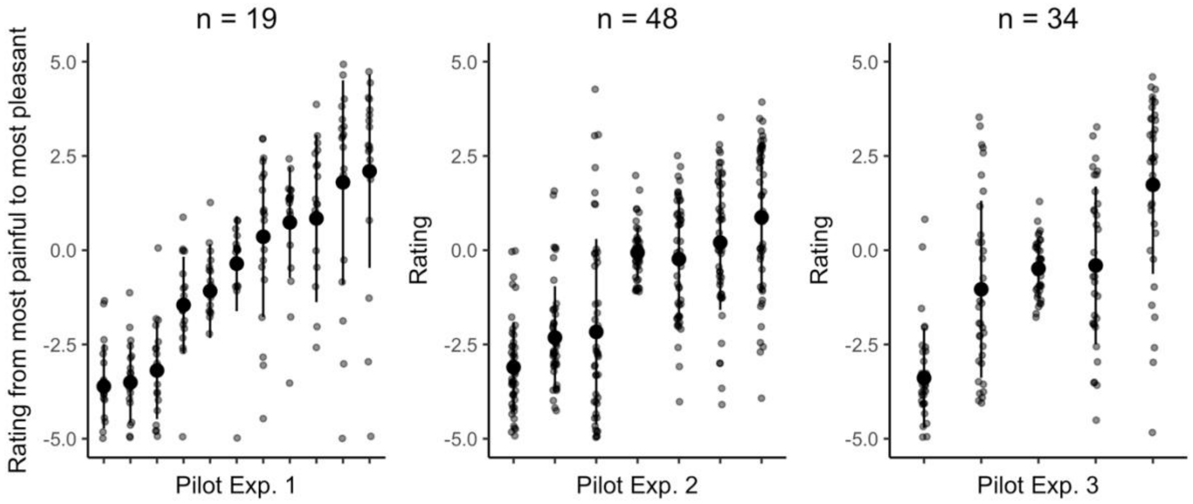


**Figure S1. Parametric shift in ratings from extreme pain to extreme pleasure for observed robotic expressions.** Average and standard deviation rating and individual rating is shown. Data from participants that did not complete the entire experiment were removed, as well as data from participants who finished the experiment too fast to pay attention to the videos (Exp. 1: *n* = 2, one additional participant indicated that he/she didn't follow instruction, Exp. 2: *n* = 5, Exp. 3: *n* = 6). Participants were instructed to rate each video on a continuum from 'most painful (-5) to 'most pleasant (+5). Videos were presented five (Exp. 1) or four (Exp. 2 and 3) times in a randomised order. In Pilot Experiments 1 and 2, participants completed the Negative Attitudes towards Robots Scale (NARS) [1,2] at the end of the experiment, as part of a different research project.

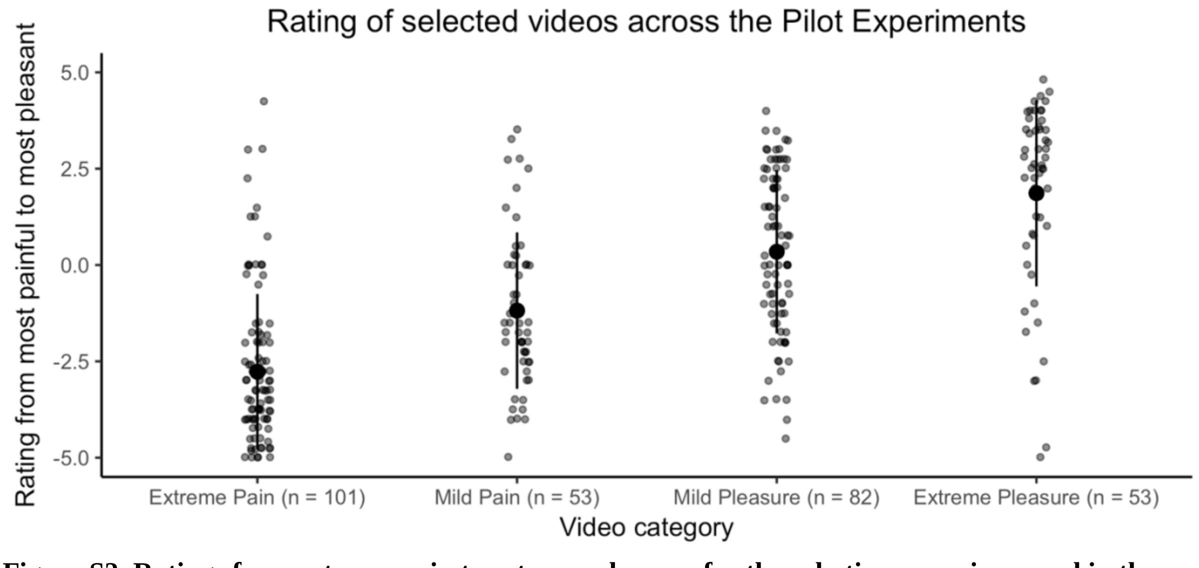


**Figure S2. Ratings from extreme pain to extreme pleasure for the robotic expressions used in the main task.** Average and standard deviation rating and individual rating is shown across the three Pilot Experiments. *n* indicates the number of participants that rated the videos across the Pilot Experiments.

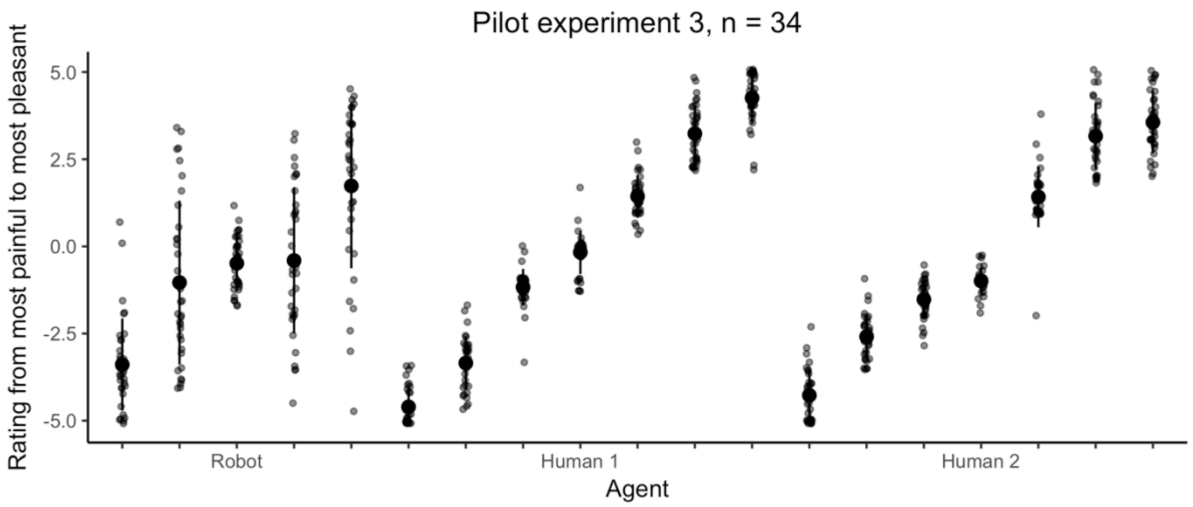


**Figure S3. Parametric shift in ratings from extreme pain to extreme pleasure for observed robotic and human expressions.** Average and standard deviation rating and individual rating is shown. Participants first rated robot videos followed by the human videos. The videos of Human 1 were used in the fMRI experiment.

**Table S1. Coordinates for parcels for the group-contrained subject-specific fROI analyses**

| *Parcel* | *MNI Coordinates* | | |
|---|---|---|---|
| | *x* | *y* | *z* |
| Anterior middle cingulate cortex | 0 | 2 | 42 |
| Left secondary somatosensory cortex | -62 | -32 | 34 |
| Right secondary somatosensory cortex | 60 | -28 | 38 |
| Left insula | -42 | -2 | -4 |
| Right insula | 42 | 6 | -6 |
| Left middle frontal gyrus | -46 | 36 | 14 |
| Right middle frontal gyrus | 50 | 42 | 12 |

Parcels are derived from [3]. A 9mm sphere is used for all parcels.

**Table S2. Outcome of the group-constrained subject-specific fROI repetition suppression analysis for the full sample (*n* = 26)**

| *ROI* | *Effect* | | |
|---|---|---|---|
| | *Agent* | *Emotion* | *Agent * Emotion* |
| Anterior middle cingulate cortex | 0.77 **0.03** [0.31/3.24] | 2.69 **0.10** [0.85/1.18] | 0.44 **0.02** [0.08/12.22] |
| Left secondary somatosensory cortex | 0.21 **0.01** [0.22/4.52] | 3.43 **0.12** [1.55/0.65] | 0.07 **0.003** [0.09/10.78] |
| Right secondary somatosensory cortex | 0.99 **0.04** [0.29/3.47] | 0.91 **0.04** [0.34/2.95] | 0.03 **0.001** [0.03/35.48] |
| Left insula | 0.69 **0.03** [0.30/3.39] | 1.16 **0.04** [0.35/2.85] | 0.04 **0.001** [0.03/36.66] |
| Right insula | 0.004 **0** [0.24/4.22] | 1.66 **0.06** [0.35/2.90] | 0.25 **0.01** [0.02/49.04] |
| Left middle frontal gyrus | 1.43 **0.05** [0.46/2.20] | 1.85 **0.07** [0.49/2.06] | 0.11 **0.004** [0.06/16.77] |
| Right middle frontal gyrus | 0.02 **0.001** [0.23/4.40] | 2.45 **0.09** [1.25/0.80] | 2.74 **0.10** [0.12/8.04] |

*F* and $\eta^2$ are reported. Factors: agent (novel or repeated) and emotion (novel or repeated). $BF_{10}/BF_{01}$ from the Bayesian Repeated Measures ANOVA with default prior scales (r scale fixed effects = 0.5) [4-6] is reported in square brackets and denotes the evidence for $H_1$ or $H_0$. The main effect model was preferred to the interaction model for all ROIs. Dependent variable: pre – post-socialising difference score in blood oxygen level-dependent response.

**Table S3. Outcome of the group-constrained subject-specific fROI repetition suppression analysis for the selected sample (*n* = 20)**

| *ROI* | *Effect* | | |
|---|---|---|---|
| | *Agent* | *Emotion* | *Agent * Emotion* |
| Anterior middle cingulate cortex | 0.15 **0.008** [0.25/3.94] | 0.95 **0.05** [0.35/2.84] | 0.87 **0.04** [0.04/28.92] |
| Left secondary somatosensory cortex | 0.16 **0.008** [0.24/4.14] | 2.96 **0.14** [1.26/0.80] | 1.13 **0.06** [0.12/8.09] |
| Right secondary somatosensory cortex | 0.25 **0.01** [0.25/4.05] | 1.94 **0.09** [0.65/1.54] | 5.91e-4 **0** [0.05/19.50] |
| Left insula | 0.50 **0.03** [0.29/3.40] | 0.63 **0.03** [0.30/3.29] | 0.04 **0.002** [0.03/36.70] |
| Right insula | 0.004 **0** [0.23/4.30] | 0.95 **0.05** [0.32/3.17] | 0.04 **0.002** [0.03/39.85] |
| Left middle frontal gyrus | 0.77 **0.04** [0.34/2.94] | 3.07 **0.14** [0.81/1.24] | 0.46 **0.023** [0.11/9.08] |
| Right middle frontal gyrus | 0.66 **0.03** [0.28/3.55] | 3.31 **0.15** [2.95/0.34] | 3.82 **0.17** [0.48/2.07] |

*F* and $\boldsymbol{\eta^2}$ are reported. Factors: agent (novel or repeated) and emotion (novel or repeated). $BF_{10}/BF_{01}$ from the Bayesian Repeated Measures ANOVA with default prior scales (r scale fixed effects = 0.5) [4-6] is reported in square brackets and denotes the evidence for $H_1$ or $H_0$. The main effect model was preferred to the interaction model for all ROIs. Dependent variable: pre – post-socialising difference score in blood oxygen level-dependent response. Participants that did not meet the preregistered cut-off of ≥2 hours total interaction time were excluded (*n* = 6).

**Table S4. Outcome of the group-constrained subject-specific fROI discrete events analysis for the full sample (*n* = 26)**

| *ROI* | *Effect* | | |
|---|---|---|---|
| | *Agent* | *Emotion* | *Agent * Emotion* |
| Anterior middle cingulate cortex | 0.05 **0.002** [0.21/4.77] | 0.05 **0.002** [0.21/4.87] | 3.68 **0.13** [0.06/18.27] |
| Left secondary somatosensory cortex | 7.41* **0.23** [8.34/0.12] | 0.003 **0.000** [0.20/4.89] | 0.73 **0.03** [0.65/1.54] |
| Right secondary somatosensory cortex | 6.01* **0.19** [11.82/0.09] | 0.06 **0.002** [0.21/4.87] | 0.005 **0.000** [0.65/1.55] |
| Left insula | 0.003 **0** [0.21/4.85] | 0.35 **0.01** [0.22/4.51] | 0.064 **0.003** [0.01/73.91] |
| Right insula | 0.006 **0** [0.21/4.80] | 0.026 **0.001** [0.21/4.84] | 0.048 **0.002** [0.01/80.29] |
| Left middle frontal gyrus | 1.26 **0.05** [0.32/3.15] | 0.12 **0.005** [0.21/4.68] | 1.69 **0.06** [0.05/21.17] |
| Right middle frontal gyrus | 0.03 **0.001** [0.21/4.86] | 0.035 **0.001** [0.23/4.40] | 0.12 **0.005** [0.01/74.62] |

*F* and $\eta^2$ are reported. Factors: agent (human or robot) and emotion (pain or pleasure). $BF_{10}/BF_{01}$ from the Bayesian Repeated Measures ANOVA with default prior scales (r scale fixed effects = 0.5) [4-6] is reported in square brackets and denotes the evidence for $H_1$ or $H_0$. The main effect model was preferred to the interaction model for all ROIs. Dependent variable: pre – post-socialising difference score in blood oxygen level-dependent response. *$p$ < .05.

**Table S5. Outcome of the group-constrained subject-specific fROI discrete events analysis for the selected sample (*n* = 20)**

| *ROI* | *Effect* | | |
|---|---|---|---|
| | *Agent* | *Emotion* | *Agent * Emotion* |
| Anterior middle cingulate cortex | 0.24 **0.01** [0.28/3.60] | 0.21 **0.01** [0.25/4.06] | 1.08 **0.05** [0.03/36.17] |
| Left secondary somatosensory cortex | 6.47* **0.25** [5.93/0.17] | 0.026 **0.001** [0.23/4.31] | 0.12 **0.006** [0.43/2.34] |
| Right secondary somatosensory cortex | 6.28* **0.25** [13.71/0.07] | 0.14 **0.007** [0.24/4.16] | 0.10 **0.005** [0.97/1.03] |
| Left insula | 0.33 **0.02** [0.28/3.61] | 0.14 **0.007** [0.24/4.20] | 0.08 **0.004** [0.02/49.26] |
| Right insula | 0.55 **0.03** [0.32/3.16] | 0.94 **0.05** [0.30/3.36] | 0.46 **0.02** [0.05/22.18] |
| Left middle frontal gyrus | 0.89 **0.05** [0.31/3.25] | 0.02 **0.001** [0.23/4.29] | 0.37 **0.02** [0.03/38.18] |
| Right middle frontal gyrus | 0.40 **0.02** [0.26/3.81] | 0.02 **0.001** [0.23/4.35] | 0.001 **0** [0.03/37.64] |

*F* and $\eta^2$ are reported. Factors: agent (human or robot) and emotion (pain or pleasure). $BF_{10}/BF_{01}$ from the Bayesian Repeated Measures ANOVA with default prior scales (r scale fixed effects = 0.5) [4-6] is reported in square brackets and denotes the evidence for $H_1$ or $H_0$. The main effect model was preferred to the interaction model for all ROIs. Dependent variable: pre – post-socialising difference score in blood oxygen level-dependent response. Participants that did not meet the preregistered cut-off of ≥2 hours total interaction time were excluded (*n* = 6). $*p < .05$.

| Table S6. Whole brain repetition suppression results (*n* = 26) | | | | | | | | | |
|---|---|---|---|---|---|---|---|---|---|
| **Anatomical region** | BA | MNI Coordinates X | Y | Z | Putative functional name | T value | Cluster size | $P_{FWE}$ corrected | $P$ uncorrected |
| **a. Main Effect of Agent (novel Agent repeated Emotion > repeated Agent repeated Emotion)** | | | | | | | | | |
| ***Session 1*** | | | | | | | | | |
| **L precuneus** | **7** | **-9** | **-73** | **37** | **SPL** | **6.21** | **412** | **<0.001** | **<0.001** |
| **R precuneus** | **7** | **15** | **-61** | **31** | | **5.42** | | | **<0.001** |
| **R precuneus** | **7** | **6** | **-67** | **37** | | **5.00** | | | **<0.001** |
| **L superior occipital gyrus** | **18** | **-9** | **-97** | **4** | **V1** | **5.39** | **68** | **0.043** | **<0.001** |
| **L calcarine gyrus** | **17** | **-3** | **-82** | **-8** | **V1** | **4.17** | | | **<0.001** |
| **L superior occipital gyrus** | **18** | **-15** | **-88** | **10** | **V1** | **4.11** | | | **<0.001** |
| **L fusiform gyrus** | **37** | **-36** | **-43** | **-23** | **FG4** | **4.88** | **76** | **0.030** | **<0.001** |
| **L fusiform gyrus** | **37** | **-39** | **-55** | **-20** | **FG4** | **4.87** | | | **<0.001** |
| **L fusiform gyrus** | **37** | **-39** | **-64** | **-17** | **FG2** | **3.58** | | | **<0.001** |
| **L angular gyrus** | **7/40** | **-45** | **-61** | **37** | **IPL** | **4.56** | **69** | **0.041** | **<0.001** |
| **L intraparietal sulcus** | **7/40** | **-33** | **-52** | **28** | **IPS** | **4.10** | | | **<0.001** |
| **L angular gyrus** | **7/40** | **-42** | **-64** | **52** | **IPL** | **3.75** | | | **<0.001** |
| R medial occipital cortex | 17 | 27 | -49 | -2 | V1 | 4.47 | 11 | 0.773 | <0.001 |
| R fusiform gyurs | 37 | 36 | -40 | -20 | FG3 | 4.43 | 40 | 0.150 | <0.001 |
| R fusiform gyrus | 37 | 39 | -52 | -20 | FG4 | 3.79 | | | <0.001 |
| L posterior fusiform gyrus | 18 | -30 | -76 | -14 | V4 | 4.27 | 10 | 0.565 | <0.001 |
| L middle frontal gyrus | 9 | -33 | 14 | 43 | MFG | 4.14 | 14 | 0.548 | <0.001 |
| L middle temporal gyrus | 21 | -45 | -37 | -5 | MTG | 4.07 | 10 | 0.803 | <0.001 |
| L anterior occipital cortex | 17 | -24 | -52 | 1 | V2 | 4.07 | 24 | 0.416 | <0.001 |

| | | | | | | | | | |
|---|---|---|---|---|---|---|---|---|---|
| L parahippocampal gyrus | 28 | -21 | -40 | -5 | | 4.01 | | | <0.001 |
| R lingual/fusiform gyrus | 19 | 30 | -70 | 1 | | 4.02 | 20 | 0.512 | <0.001 |
| R anterior occipital cortex | 18 | 36 | -61 | 1 | | 4.01 | | | <0.001 |
| R superior occipital gyrus | 19 | 24 | -76 | 4 | V1 | 3.62 | | | 0.001 |
| ***Session 2*** | | | | | | | | | |
| L medial temporal lobe | 37/39 | -39 | -46 | 4 | | 5.34 | 48 | 0.119 | <0.001 |
| L hippocampus/dentate gyrus | | -24 | -40 | 1 | | 3.57 | | | <0.001 |
| L thalamus | | -30 | -31 | 4 | | 3.48 | | | <0.001 |
| L precuneus | 7 | -12 | -67 | 34 | SPL | 4.91 | 10 | 0.806 | <0.001 |
| R hippocampus/dentate gyrus | | 27 | -37 | 1 | | 4.54 | 24 | 0.420 | <0.001 |
| L fusiform gyrus | 37 | -39 | -58 | -20 | FG4 | 4.40 | 26 | 0.378 | <0.001 |
| L inferior temporal gyrus | 37 | -54 | -52 | -17 | ITG | 4.13 | | | <0.001 |
| R fusiform gyrus | 37 | 42 | -52 | -20 | FG4 | 4.08 | 17 | 0.597 | <0.001 |
| **b. Main Effect of Emotion (rAnE > rArE)** | | | | | | | | | |
| ***Session 1*** | | | | | | | | | |
| R middle temporal gyrus/inferior parietal lobule | 40 | 57 | -40 | 7 | MTG/IPL | 4.40 | 36 | 0.186 | <0.001 |
| ***Session 2*** | | | | | | | | | |
| *No suprathreshold clusters survived this contrast* | | | | | | | | | |

All contrasts were evaluated at p < 0.001, k = 10 voxels, uncorrected. Regions that survived cluster correction at p < 0.05 FWE-corrected are denoted by bold font. Clusters with three local maxima greater than 8.00mm apart are listed.

| Table S7. Discrete coded events design: Main effects of agent and interaction between agent and scan session (*n* = 26) | | | | | | | | | |
|---|---|---|---|---|---|---|---|---|---|
| **Anatomical region** | BA | MNI Coordinates | | | Putative functional name | T value | Cluster size | $P_{FWE}$ corrected | *P* uncorrected |
| | | X | Y | Z | | | | | |
| **a. Main Effect of Agent: Human > Robot** | | | | | | | | | |
| ***Session 1*** | | | | | | | | | |
| **L precuneus** | **7** | **-9** | **-73** | **37** | **SPL** | **6.21** | **412** | **<0.001** | **<0.001** |
| **R precuneus** | **7** | **15** | **-61** | **31** | | **5.42** | | | **<0.001** |
| **R precuneus** | **7** | **6** | **-67** | **37** | **SPL** | **5.00** | | | **<0.001** |
| ***Session 2*** | | | | | | | | | |
| **R calcarine gyrus** | **18** | **12** | **-88** | **13** | **V2** | **10.50** | **208** | **<0.001** | **<0.001** |
| **L cuneus** | **19** | **-3** | **-91** | **16** | **V3d** | **9.62** | | | **<0.001** |
| **L calcarine cortex** | **18** | **-3** | **-94** | **1** | **V1** | **8.74** | | | **<0.001** |
| **L ventral occipital cortex** | **19** | **-9** | **-76** | **-14** | **V3v** | **8.58** | **26** | **<0.001** | **<0.001** |
| **R lingual gyrus** | **17** | **12** | **-64** | **-2** | **V2** | **6.70** | **14** | **<0.001** | **<0.001** |
| **R calcarine gyrus** | **17** | **18** | **-64** | **4** | **V1** | **6.41** | | | **<0.001** |
| **L calcarine gyrus** | **17** | **-3** | **-70** | **10** | **V1** | **6.39** | **10** | **0.001** | **<0.001** |
| **L calcarine gyrus** | **18** | **-6** | **-79** | **7** | **V1** | **6.10** | | | **<0.001** |
| **b. Main Effect of Agent: Robot > Human** | | | | | | | | | |
| ***Session 1*** | | | | | | | | | |
| **R middle occipital gyurus** | **19** | **39** | **-82** | **13** | **MOG** | **15.97** | **430** | **<0.001** | **<0.001** |
| **R angular gyrus** | **7** | **30** | **-61** | **46** | **IPS** | **9.99** | | | **<0.001** |
| **R inferior occipital gyrus** | **19** | **39** | **-85** | **-2** | **V4** | **9.75** | | | **<0.001** |
| **L inferior occipital gyrus** | **37/19** | **-45** | **-61** | **-11** | **FG4** | **14.71** | **221** | **<0.001** | **<0.001** |

| | | | | | | | | | |
|---|---|---|---|---|---|---|---|---|---|
| **L fusiform gyrus** | **37** | **-30** | **-61** | **-11** | **FG3** | **12.52** | | | **<0.001** |
| **R fusiform gyrus** | **37** | **27** | **-58** | **-14** | **FG3** | **14.61** | **272** | **<0.001** | **<0.001** |
| **R ventral temporal cortex** | **37** | **36** | **-58** | **-8** | **FG1** | **10.10** | | | **<0.001** |
| **R inferior temporal gyrus** | **37** | **45** | **-61** | **-8** | **FG4** | **10.08** | | | **<0.001** |
| **L middle occipital gyrus** | **19** | **-36** | **-85** | **10** | **MOG** | **10.15** | **175** | **<0.001** | **<0.001** |
| **L middle occipital gyrus** | **19** | **-30** | **-82** | **4** | **MOG** | **6.72** | | | **<0.001** |
| **L middle occipital gyrus** | **19** | **-30** | **-76** | **19** | **MOG** | **7.34** | | | **<0.001** |
| **L superior parietal lobule** | **7** | **-27** | **-64** | **46** | **SPL** | **8.97** | **105** | **<0.001** | **<0.001** |
| **L inferior parietal lobule** | **40** | **-36** | **-58** | **52** | **IPL/IPS** | **7.88** | | | **<0.001** |
| **L angular gyrus** | **7/40** | **-33** | **-52** | **37** | **IPS** | **7.16** | | | **<0.001** |
| **R precentral gyrus** | **44** | **51** | **11** | **31** | **PMv** | **8.35** | **62** | **<0.001** | **<0.001** |
| **R precentral gyrus** | **44** | **45** | **5** | **28** | | **7.02** | | | **<0.001** |
| **L inferior frontal gyrus** | **45** | **-27** | **23** | **1** | **IFG** | **7.68** | **15** | **<0.001** | **<0.001** |
| **R IFG, pars triangularis** | **45** | **48** | **35** | **13** | **IFG** | **6.56** | **16** | **<0.001** | **<0.001** |
| **R middle frontal gyrus** | **45** | **45** | **32** | **22** | **MFG** | **6.45** | | | **<0.001** |
| ***Session 2*** | | | | | | | | | |
| **R middle occipital gyrus** | **19** | **39** | **-79** | **13** | **MOG** | **15.23** | **179** | **<0.001** | **<0.001** |
| **R inferior occipital gyrus** | **19** | **36** | **-82** | **-2** | **IOG** | **10.15** | | | **<0.001** |
| **R fusiform gyrus** | **37** | **27** | **-58** | **-14** | **FG3** | **13.25** | **112** | **<0.001** | **<0.001** |
| **L middle occipital gyrus** | **19** | **-33** | **-85** | **13** | **MOG** | **11.20** | **179** | **<0.001** | **<0.001** |
| **L middle occipital gyrus** | **19** | **-33** | **-82** | **1** | **MOG** | **9.31** | | | **<0.001** |
| **L fusiform gyrus** | **37** | **-30** | **-61** | **-11** | **FG3** | **10.84** | **158** | **<0.001** | **<0.001** |
| **L inferior occipital gyrus** | **19** | **-45** | **-64** | **-11** | **FG4** | **10.34** | | | **<0.001** |
| **R inferior temporal gyrus** | **37** | **51** | **-61** | **-14** | **FG2** | **8.88** | **32** | **<0.001** | **<0.001** |
| **R inferior temporal gyrus** | **37** | **45** | **-61** | **-8** | **FG4** | **7.45** | | | **<0.001** |
| **R superior parietal lobule** | **7** | **24** | **-73** | **55** | **SPL** | **8.75** | **148** | **<0.001** | **<0.001** |

| | | | | | | | | | |
|---|---|---|---|---|---|---|---|---|---|
| **R angular gyrus** | **7** | **30** | **-58** | **49** | **IPS** | **8.33** | | | **<0.001** |
| **R middle occipital gyrus** | **39** | **30** | **-73** | **34** | **MOG** | **7.47** | | | **<0.001** |
| **R precentral gyrus** | **45** | **48** | **8** | **34** | **PMv** | **8.15** | **34** | **<0.001** | **<0.001** |
| **L inferior parietal lobule** | **40** | **-27** | **-58** | **43** | **IPL** | **7.50** | **41** | **<0.001** | **<0.001** |
| **L middle frontal gyrus** | **9/6** | **-36** | **-1** | **31** | **MFG** | **6.81** | **13** | **<0.001** | **<0.001** |
| c. Agent by Scan Session Interactions | | | | | | | | | |
| ***Robot > Human, Session 1 > Session 2*** | | | | | | | | | |
| **L superior parietal lobule** | **7** | **-24** | **-55** | **52** | **SPL/IPS** | **5.10** | **113** | **0.005** | **<0.001** |
| **L superior parietal lobule** | **7** | **-33** | **-49** | **55** | **SPL** | | | | **<0.001** |
| **L intraparietal sulcus** | **7/40** | **-39** | **-43** | **49** | **IPS** | | | | **<0.001** |
| ***Robot > Human, Session 2 > Session 1*** | | | | | | | | | |
| *No suprathreshold clusters emerge* | | | | | | | | | |

All contrasts were evaluated at $p < 0.05$, k = 10 voxels, FWE-corrected. Regions that survived cluster correction at $p < 0.05$ FWE-corrected are denoted by bold font. Clusters with three local maxima greater than 8.00mm apart are listed.

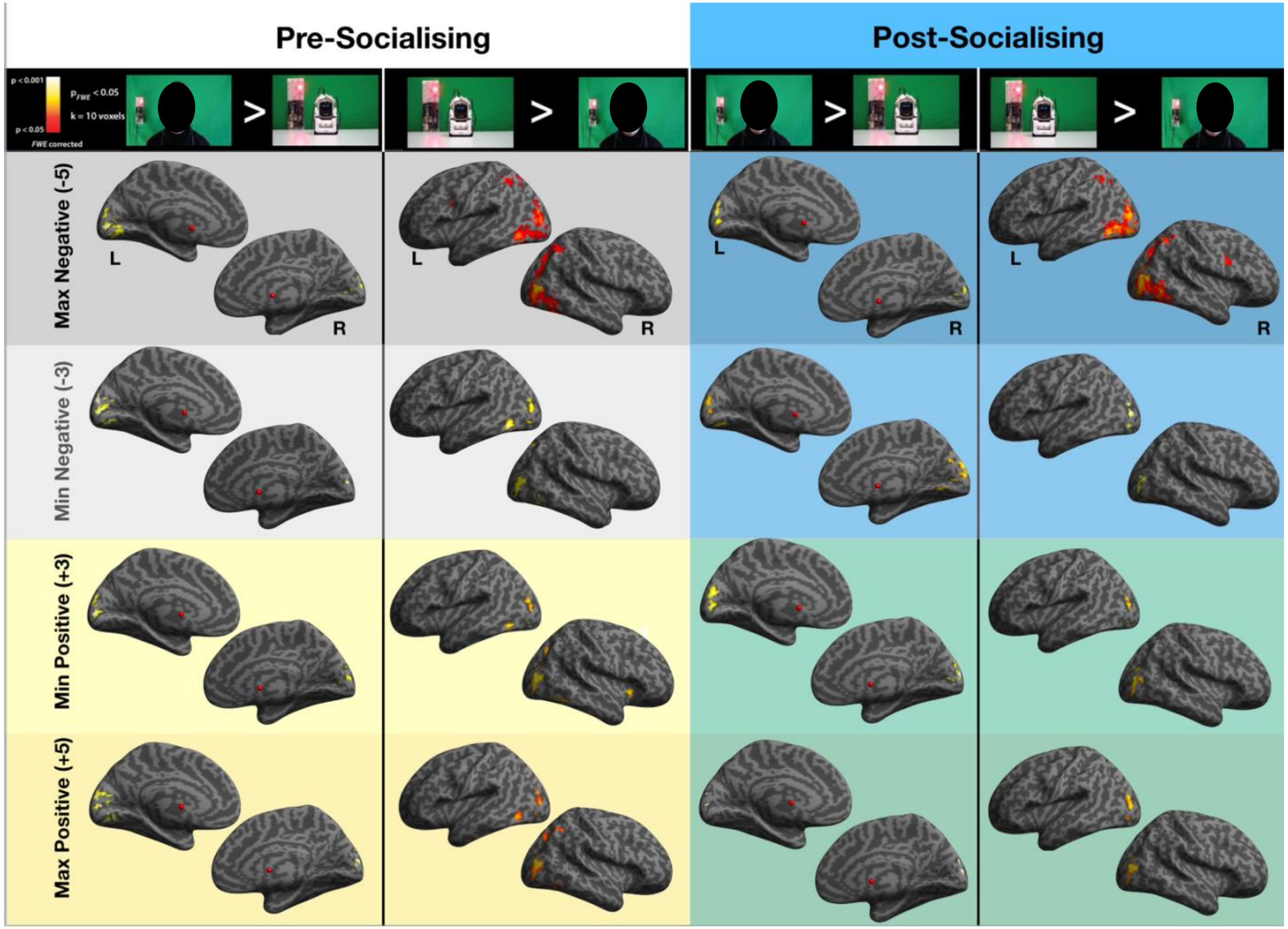


**Figure S4. Visualisations of the main effect of watching videos of a human compared to a robot (and vice versa) during the pre-socialising and post-socialising scan sessions, broken down by emotional valence and intensity.** Each column represents a single contrast comparing the perception of the two agents, with the pre-socialising contrasts visualised in the left two columns and the post-socialising contrasts visualised in the right two columns. Each row represents the valence and intensity of the observed video, ranging from the most negative videos (top row) to the most positive videos (bottom row). All contrasts were evaluated at the p(FWE) < 0.05 and k = 10 voxels threshold. Please note that images of the human agent have been obscured due to bioRxiv's policy to not post manuscript that contain identifying material.
